# *TRA1*: a locus responsible for controlling *Agrobacterium*-mediated transformability in barley

**DOI:** 10.1101/2019.12.19.882274

**Authors:** Beata Orman-Ligeza, Wendy Harwood, Pete E. Hedley, Alison Hinchcliffe, Malcolm Macaulay, Cristobal Uauy, Kay Trafford

**Author notes:** Correspondence: Kay Trafford.

## Abstract

In barley (*Hordeum vulgare* L.), *Agrobacterium*-mediated transformation efficiency is highly dependent on genotype with very few cultivars being amenable to transformation. Golden Promise is the cultivar most widely used for barley transformation and developing embryos are the most common donor tissue. We tested whether barley mutants with abnormally large embryos were more or less amenable to transformation and discovered that mutant M1460 had a transformation efficiencies similar to that of Golden Promise. The large-embryo phenotype of M1460 is due to mutation at the *LYS3* locus. There are three other barley lines with independent mutations at the same *LYS3* locus, and one of these, Risø1508 has an identical missense mutation to that in M1460. However, none of the *lys3* mutants except M1460 were transformable showing that the locus responsible for transformation efficiency, *TRA1*, was not *LYS3* but another locus unique to M1460. To identify *TRA1*, we generated a mapping population by crossing M1460 to the cultivar Optic, which is recalcitrant to transformation. After four rounds of backcrossing to Optic, plants were genotyped and their progeny were tested for transformability. Some of the progeny lines were transformable at high efficiencies similar to those seen for the parent M1460 and some were not transformable, like Optic. A region on chromosome 2H inherited from M1460 is present in transformable lines only. We propose that one of the 225 genes in this region is *TRA1*.

## INTRODUCTION

*Agrobacterium*-mediated transformation of immature barley embryos was first reported in 1997 using the cultivar Golden Promise (Tingay *et al*., 1997) and most barley transformation since then has used this cultivar. This is because no other cultivar has been found to give a similar or higher transformation efficiency (Bartlett *et al*., 2008; Harwood *et al*., 2009; Harwood, 2012). Wheat, like barley, is also transformable using *Agrobacterium* but despite successful attempts to increase transformation efficiency (Hayta *et al*., 2019), the process is still cultivar dependant: cv Fielder being one of the most responsive cultivars. Efficiencies as high as 86.7% can be obtained with barley cv Golden Promise when acetosyringone, which triggers the transformation activity of *Agrobacterium*, is added to the co-cultivation media (Hensel *et al*., 2008). However, Golden Promise has certain drawbacks: it is an old cultivar first developed in 1956 by mutagenesis of the cultivar Maythorpe (Forster, 2001), it has a low yield compared with modern elite barley cultivars and it is prone to disease, particularly mildew (Douchkov *et al*., 2014).

The gene(s) underlying transformability in Golden Promise are not known, but recent studies indicated that three genomic regions are required. One region, TFA1 is located on chromosome 3H and two others, TFA2 and TFA3, are both on chromosome 2H (Hisano and Sato, 2016; Hisano *et al*., 2017). This suggests that the transformability trait in Golden Promise a multi-genic in nature which means that it would be technically challenging to transfer the trait to another barley cultivar. With the increasing public acceptance of genetically modified crops on the horizon, we anticipate a need to transform elite barley cultivars directly rather than transferring transgenes from Golden Promise to elite lines by crossing. Also, for academic as well as commercial purposes, it would be advantageous to be able to compare transgene constructs in different genetic backgrounds. Thus, there is a need for other, preferably elite, lines of barley that can be transformed with high efficiency.

Testing immature barley embryos from several elite cultivars has shown that they do not form callus in tissue culture as readily as Golden Promise (for examples see Zalewski *et al*., 2012; Murray *et al*., 2004; Hensel *et al*., 2008; Cho *et al*., 1998). This suggests that the ability of the cultured embryos of Golden Promise to form callus is unusual and this may contribute to its unique transformability. The plant phytohormone auxin is known to promote cell division in un-differentiated cells (Hiei *et al*., 2014) and to be synthesised by plant cells upon infection by *Agrobacterium* (Gohlke *et al*., 2014). The developing embryos of Golden Promise have higher levels of auxin (and also lower levels of salicylic and abscisic acid) than embryos of non-transformable cultivars (Hisano *et al*., 2016). This suggests that unusual phytohormone levels may be required for high transformation efficiency.

Whilst working on mutants of barley affected at the *LYS3* locus, we noticed that their developing embryos had an unusual cellular organization reminiscent of callus tissue (Cook *et al*., 2018). The *LYS3* gene is a transcription factor called Prolamin Binding Factor, and is expressed in developing and germination grain (Moehs *et al*., 2019; Orman-Ligeza *et al*., 2019. The scutellar epidermis in *lys3* mutants consists of multiple layers of cells instead of the usual single layer (Deggerdal *et al*., 1986; Cook *et al*., 2018). As well as abnormal cellular structure, *LYS3* mutant embryos are larger than normal and the grains show a range of other unusual phenotypes, including high lysine content, shrivelled seeds and low starch content (Doll, 1974; Talberg, 1973; Trafford and Fincher, 2014; Cook *et al*., 2018).

Here, we investigate the transformation efficiency of the *lys3* mutant lines compared with that of their wild-type parents, and Golden Promise. We found that one mutant, M1460, but not three other *lys3* mutants, is amenable to transformation. To discover the genetic basis of this transformability trait, we generated a population of plants segregating for transformability by crossing M1460 and the elite, non-transformable cultivar Optic. We used this population to identify a locus, *<u>TRA1</u>* responsible for <u>tra</u>nsformability in barley.

## MATERIALS AND METHODS

### Barley Germplasm

The collection of barley mutants and wild-type Minerva used here was requested from the following publicly-available germplasm collections: BBSRC Cereals Collection, JIC (Bomi, Optic, Maythorpe, Golden Promise, Risø1508 and Minerva Accessions 3524, 7732 and 9562), USDA-ARS National Small Grains Collection, Idaho, USA (CIho10086, CIho11477, PI243181, PI247923, PI321813 and PI328951), and Nordic Gene Bank, Alnarp, Norway (B9609). Minerva, Risø18, Risø19, and M1460 were a kind gift from Birthe Møller Jespersen, University of Copenhagen, Denmark.

### Plant Growth

For transformation experiments, plants were grown in a controlled environment room with a 16-h day length, maintained at 15 °C day and 12 °C night temperatures, 80% relative humidity and with light levels of 500 μmol.m^−2^.s^−1^ at the mature plant canopy level provided by metal halide lamps (HQI) supplemented with tungsten bulbs.

For determination of embryo dry weight and for genotyping, individual grains were germinated in Petri dishes on moist filter paper, incubated at 4 °C overnight and then transferred to room temperature. When roots and shoots were established, each seedling was transplanted into a 1 L pot containing Levington M2 compost (Everris, Geldermalsen, The Netherlands) and grown in a glasshouse maintained at 15 °C (night) and 20 °C (day) with additional lighting provided by sodium lamps to give a minimum of 16 h of light per day.

### Measurement of Embryo Weight

Mature grains were imbibed in water overnight and then the embryos were excised. The embryo and non-embryo samples were separately oven-dried for 48 h at 65 °C and weighed.

### Testing Transformation Efficiency

Barley was transformed with *35S:GUS* using the method described in Hinchliffe and Harwood *et al*. (2019). In brief, the protocol involves use of *Agrobacterium tumefaciens* strain AGL1 and the transformation vector pBract204. This vector contains a selectable marker gene, *hygromycin phosphotransferase* (*HPT*) that confers hygromycin resistance (driven by a 35S promoter) and a reporter gene, *β-glucuronidase* (*GUS*) (driven by a maize (*Zea mays* L.) ubiquitin promoter). Three different basic plant tissue culture media were used during the transformation and regeneration process: callus induction, transition and regeneration media (Hinchliffe and Harwood *et al*. (2019). All media contained hygromycin at 50 mg/l unless otherwise stated.

Immature embryos (normal size) were harvested when the embryos were 1.5-2.0 mm in diameter, as described in Hinchliffe and Harwood (2019). At this stage, the grains were plump but the endosperm was still liquid. For the large-embryo mutant barley lines, embryos were harvested at a visibly similar developmental stage, even if their diameter was larger than 1.5-2.0 mm.

Following inoculation of immature embryos with *Agrobacterium*, embryos were placed scutellum side up on callus induction medium but with no hygromycin. After three days co-cultivation, the embryos were transferred to plates containing callus induction medium with hygromycin. After six weeks selection on callus induction medium (with two transfers to fresh callus induction plates), the embryo-derived callus was transferred to transition medium. After two weeks, transformed calli started to produce green areas and small shoots and the regenerating calli were transferred to regeneration medium. When shoots were 2-3 cm in length and roots had formed, the small plantlets were removed from the plates and transferred to glass culture tubes containing callus induction medium but without growth regulators. Once rooted plants reached the top of the tubes they were transferred to soil. Leaf samples were collected once the plants were established in soil and analysed to confirm the presence of the introduced genes. Transformation efficiency was calculated as the number of independent transformed lines obtained as a percentage of the number of embryos cultured.

DNA was extracted from leaf samples using the method of Fulton *et al*. (1995) and the concentration was adjusted to 10 ng/μl. The PCR reaction (in a final volume of 20 μl) contained: 1 μl of genomic DNA template, 0.5 μl of 10 mM forward primer, 0.5 μl of 10 mM reverse primer, 0.6 μl of 10 mM dNTPs, 2 μl of 10 x PCR buffer, 0.2 μl of FastStart Taq DNA polymerase (Roche, 04659163103) and water. The PCR conditions were: 95 °C for 3 min, 38 cycles of (95 °C for 30 s, 58 °C for 30 s and 72 °C for 1 min) and then 72 °C for 5 min. For *GUS* amplification, the amplicon size was 710 bp and the primers were *DR48: GUS in pBRACT204_F* (TAGATATCACACTCTGTCTG) and *DR37: GUS in pBRACT204_R* (GGAATTGATCAGCGTTGGTG). For *HYG* amplification, the amplicon size was 1035 bp and the primers were *Hygromycin_F (*TAGGAGGGCGTGGATATGT) and *Hygromycin_R* (TACACAGCCATCGGTCCAGA).

### Testing Regeneration Efficiency

This was performed as for transformation testing, except that the *Agrobacterium*-inoculation and co-cultivation steps were omitted and all media lacked hygromycin. The number of regenerated shoots (greater than 4 mm in length) from the callus regenerated from 50 individual immature embryos of each genotype was recorded.

### Measurement of Callus Size

After four to five weeks (for regeneration tests) and seven to eight weeks (for transformability tests), Petri dishes were photographed and the areas of calli were measured using ImageJ (https://imagej.nih.gov/; version 1.43u).

### Generation of the *TRA1* Mapping Population

To generate a population of plants segregating for transformation efficiency (and for mutation at the *lys3* locus that confers a large-embryo phenotype), we crossed the *lys3* mutant M1460 to the elite non-transformable malting barley cultivar, Optic. An F_1_ plant resulting from this primary cross was crossed again to Optic and the backcross 1 (BC_1_) F_1_ plants were allowed to self-pollinate to give BC_1_ F_2_ grains. The F_2_ grains were screened visually and those with large embryos (indicating that they were homozygous *lys3* mutants) were planted and allowed to self-pollinate. The large-embryo phenotype was checked again in the BC_1_F_3_ and subsequent generations (progeny testing). Developing embryos from some of these BC_1_-derived plants were used for transformation trials.

One of the BC_1_F_3_ plants was backcrossed to Optic. BC_2_ lines were generated from this cross as for BC_1_ lines above except that the large-embryo phenotype of the BC_2_-derived grains was not confirmed by progeny testing. Some BC_2_-derived plants were used for transformation trials and others were used for further backcrossing. BC_3_ and BC_4_ lines were generated as for BC_2_.

### Barley Genotyping and Data Analysis

DNA was extracted using DNeasy Plant Mini Kit (Qiagen, UK) following the manufacturer’s instructions. DNA was quantified and quality checked using a spectrometer (NanoDrop 1000, Thermo Scientific). DNA with absorbance ratios at both 260/280nm and 260/230nm of >1.8 was used and the DNA concentration was adjusted to 20 ng/μl. A total amount of 300 ng DNA per sample were lyophilized and sent to Geneseek (Neogen Europe, Ltd., Auchincruive, UK) for Illumina HTS processing and HiScan array imaging (Illumina, San Diego, CA, USA). R and Theta scores were extracted from resulting idat files using GenomeStudio Genotyping Module v2.0.2 (Illumina, San Diego, CA, United States) and allele scores were created using paRsnps (an in-house software package for clustering, visualising and comparing Illumina SNP genotyping data).

All SNP datasets in this study were analysed using R software (v.2.15.0, www.r-project.org) as follows. The genetic relationship between Minerva accessions was visualised by generating an unrooted phylogenetic tree constructed by the neighbour-joining method (with Euclidean distance matrix setting) and a Principal Coordinate Analysis (PCoA). To delimit the genomic region responsible for M1460 transformability, alleles of M140 and the recurrent parent Optic were color-coded and visualized using the heatmap.2 package within R (Warnes *et al*., 2015). The distribution of polymorphic markers across all seven barley chromosomes was visualized using ggplot2 package within R (Wickham, 2016).

### Candidate Genes in the *TRA1* Region

The Gene Ontologies (GOs) of the 225 genes in the *TRA1* region were downloaded from Ensembl Plants using BioMart (https://plants.ensembl.org/) and were analysed using a GO terms categoriser (www.animalgenome.org/cgi-bin/util/gotreei) with default parameters and the ‘plant GO slim’ option (Hu *et al*., 2008). GOs were then analysed using Revigo (http://revigo.irb.hr/revigo.jsp) to find a representative subset of GO terms and to reduce and visualize these. Bubble size indicates the frequency of the GO term in the GO database, Revigo. Highly similar GO terms are linked by edges in the graph (Supek *et al*., 2011).

## RESULTS

### The *lys3* Mutant, M1460 is Amenable to Transformation

Following chemical mutagenesis, four *lys3* mutant lines were identified: Risø1508, Risø18 and Risø19 (derived from the parent cultivar Bomi; Tallberg,1973; Munck,1992) and M1460 (derived from Minerva; Aastrup,1983). As described previously, *lys3* embryos are larger than normal (Cook *et al*., 2018; Fig. 1), particularly those of M1460 (325% of fresh weight of the wild type cultivar Minerva; Fig. S1). For comparison, we measured the size of the mature embryos of Golden Promise and Maythorpe and both genotypes have normal-sized embryos, similar to Minerva (Fig. S1). However, we also observed cell proliferation around the edge of the scutellum in the immature embryos of M1460. This is normally only observed in embryos following culture on media containing high levels of auxin (Fig. 1). This led us to test whether *lys3* influences the performance of the embryos in tissue culture.

**Figure 1.**
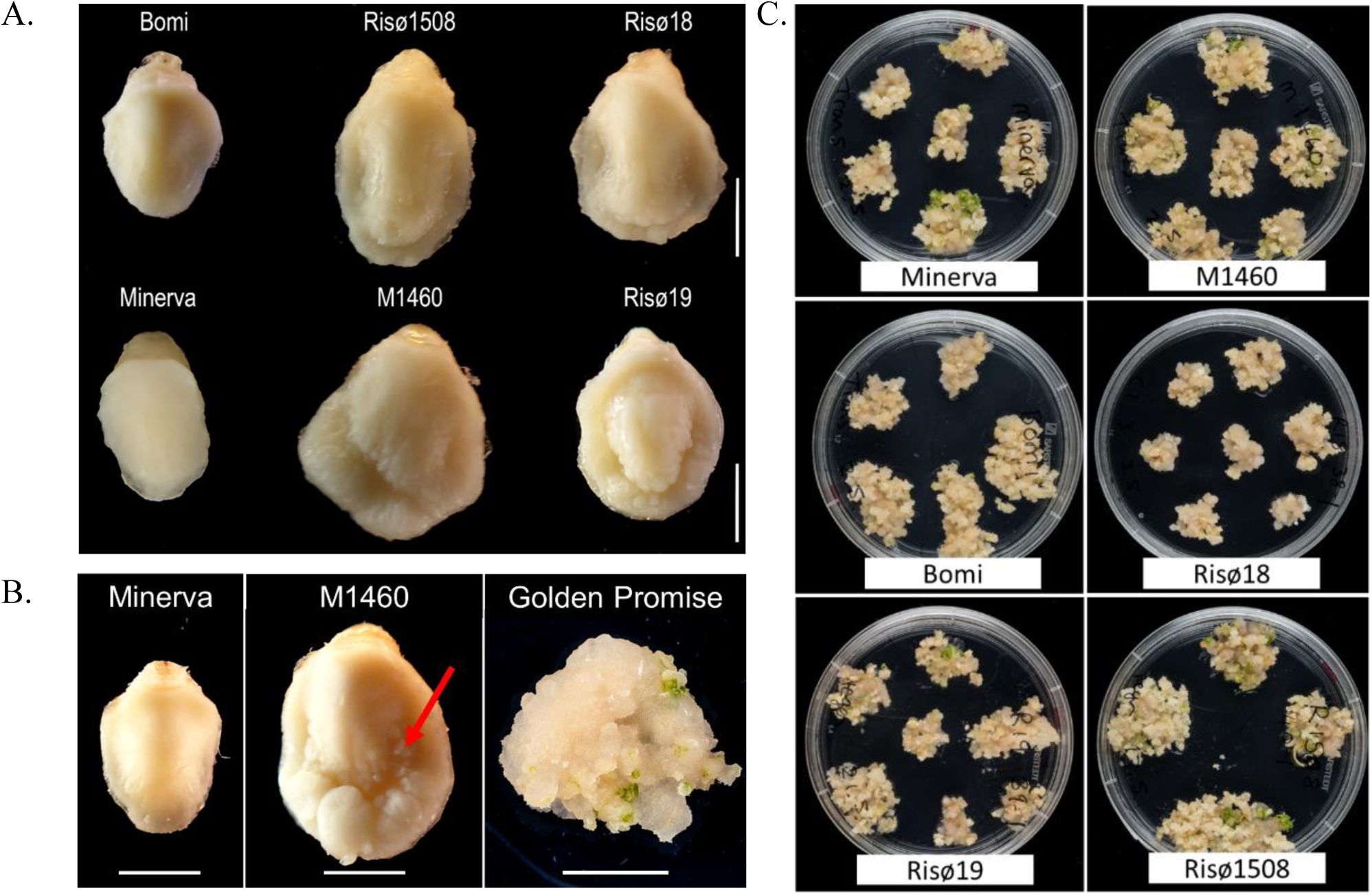
Embryo morphology and callus formation. A. Embryos from mature grains. Bomi and Minerva are wild-type controls and Risø18, Risø19, Risø1508 and M1460 are *lys3* mutants. Bar = 2 mm. B. Immature barley embryos excised from developing grains and a typical Golden Promise embryo after growth on callus induction and transition media. The arrow indicates cell proliferation around the edge of the scutellum of the large-embryo mutant, M1460. Bars = 2 mm for Minerva and M1460 embryos and xx mm for the Golden Promise callus. C. Representative plates showing individual embryos after growth on callus induction and transition media.

To investigate the responses of immature *lys3* embryos in tissue culture (in the absence of *Agrobacterium* co-cultivation or selection on hygromycin), developing embryos were grown on culture induction medium and then transferred to regeneration medium to induce shoots. All genotypes tested developed some callus and produced shoots. No consistent genotype-dependent differences were detected and there was no correlation between embryo size and callus size, suggesting that the large-embryo trait *per se* does not affect callus induction. The extent of callus growth and the number of shoots produced per embryo varied between replicates, lines and experiments.

To test transformation efficiency, embryos of different genotypes were transformed with *35S:GUS* and cultured on media containing hygromycin. Transformation efficiency was calculated as the number of independent transformed lines obtained as a percentage of the number of embryos cultured. Across two independent transformation experiments each involving at least 50 embryos per genotype, M1460, Golden Promise and the parent of Golden Promise, cv. Maythorpe, were all transformable but the parent of M1460, cv. Minerva failed to produce any transformants (Fig. 2). In this experiment, the mean transformation efficiency of M1460 at 13%, was lower than that of either Golden Promise or Maythorpe. However, 13% transformation efficiency is higher than all values previously recorded in our lab for cultivars other than Golden Promise or Maythorpe (Harwood, 2012).

**Figure 2.**
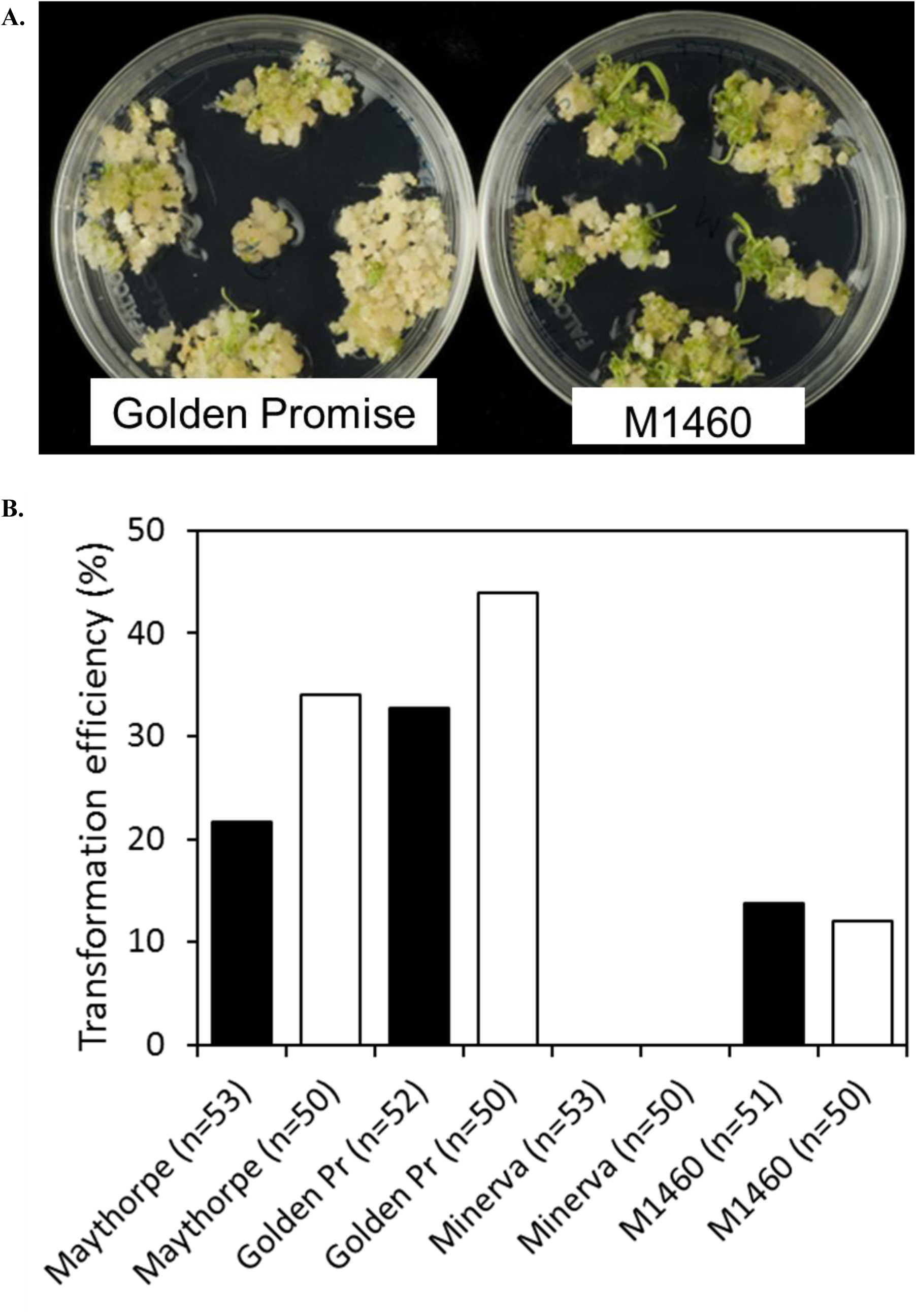
Transformation efficiency. A. Immature barley embryos were transformed with *35:GUS*. Following transformation and callus induction, calli were transferred to transition medium and then to regeneration medium. Representative plates of M1460 and Golden Promise are shown. B. Transformation efficiency was calculated as the number of independent transformed lines obtained as a percentage of the number of embryos cultured (n = number of embryos). Values for two independent experiments are shown: experiment 1 (black bars) and experiment 2 (white bars). The value for Minerva in both experiments was zero (no plants were recovered). Golden Pr = Golden Promise.

Similar transformation experiments with the other *lys3* mutants all failed (i.e. did not produce transformants). The number of embryos tested were 275 (Risø1508), 200 (Risø18) and 125 (Risø19). These data suggest that M1460 has unique properties not possessed by other *lys3* mutants that allow transformation by *Agrobacterium*. The transformability of M1460 is therefore not caused by mutation at the *lys3* locus, but by another genetic factor.

### A Region of Chromosome 2H is Responsible for M1460 Transformability

To discover the gene(s) responsible for transformability in M1460, we generated a mapping population of plants segregating for this trait by crossing Optic (a non-transformable cultivar) to M1460 (transformable). Four rounds of backcrossing to Optic generated BC_1_, BC_2_, BC_3_ and BC_4_ lines. To select homozygous *lys3* mutant lines from the F_2_ grains in the BC_1_ generation, we grew only grains that were shrunken (a characteristic phenotype of the *lys3* mutant grains).

Immature embryos from plants from each BC generation were tested for transformability with *Agrobacterium* and a *35S:GUS* vector. After co-cultivation with *Agrobacterium*, the embryos were grown on callus induction medium containing hygromycin. At this stage, we noticed that the embryos of some genotypes grew better and produced more callus than others (Fig. 3A). Embryos of the *LYS3* wild-type parents, Bomi and Minerva, and from the BC_3_ and BC_4_ lines did not grow as much as those of Golden Promise, M1460, BC_1_ and BC_2_. These differences between genotypes were also apparent when the sizes of the calli were measured (Fig. 3B). Thus, the growth of calli during transformation experiments varied with genotype. This result is different from that obtained for regeneration experiments: embryos of all genotypes grew and produced callus to a similar extent when they were grown on callus induction medium without exposure to *Agrobacterium* and with no hygromycin selection (Fig. 1C and Fig. S1B).

**Figure 3.**
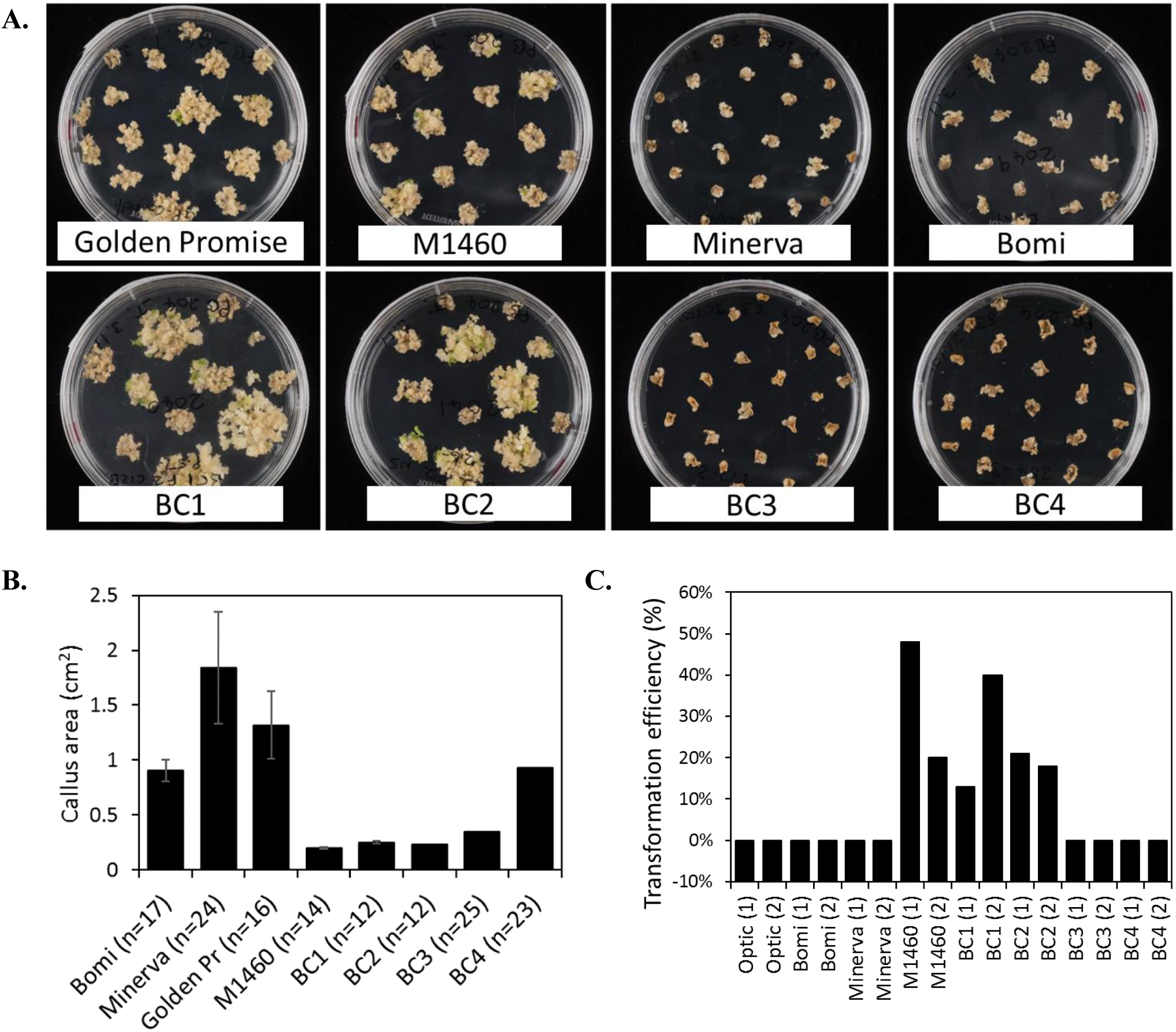
Transformation of lines derived from a cross between M1460 and Optic. Immature F_3_ or F_4_ embryos from one or two plants (P1 and P2) from each backcrossed (BC) line were tested for transformability. The embryo phenotype was visually assessed as normal-embryo (NE) or large-embryo (LE; i.e. *lys3* mutant). A. Immature barley embryos were transformed with *35:GUS*. Following transformation and callus induction, calli were transferred to transition media. Representative plates are shown. B. The area of individual calli were measured using ImageJ. Values are means ± SE for one plate (n = number of calli) from a single experiment. Golden Pr = Golden Promise. C. Transformation efficiency was calculated as the number of independent transformed lines obtained as a percentage of the number of embryos cultured. Values of zero mean that no transgenic plants were recovered. Fifty embryos were cultured per line in each experiment. Values for two independent experiments are shown: experiment 1 (1) and experiment 2 (2).

After transfer to transition medium and then to regeneration medium, plantlets and then rooted plants were regenerated from M1460, BC_1_ and BC_2_ calli. Transgenes were detected in all regenerated plants using PCR and primers specific to hygromycin and GUS (Fig. S2). The transformation efficiencies for these lines (the number of independent transformed lines obtained as a percentage of the number of embryos cultured) were comparable with those normally obtained in our lab with Golden Promise (Fig. 3C). In contrast, no shoots or regenerated plants were obtained from Optic, Bomi, Minerva, BC_3_ or BC_4_ (TE = zero, 50 embryos per line were cultured in each of two experiments) showing that these lines were not transformable.

To identify the region(s) of the M1460 genome responsible for inherited transformability, we genotyped the parents of the backcrossed population, M1460 and Optic, using the Barley 50k iSelect SNP array (Supplementary File S1). A total of 44,040 markers were evaluated of which 9,456 markers were polymorphic between Optic and M1460. The following markers were eliminated from further analysis: 31,925 markers that were monomorphic, 2,535 that gave values for one parent only, 45 that were not mapped to any chromosome, and 79 that were heterozygous in Optic or M1460. The distribution of markers was not balanced across all seven barley chromosomes: chromosome 5H was over-represented (Fig. S3).

We also genotyped the non-transformable BC_3-4_ embryo-donor plants and all the transgenic plants that were recovered from the transformation testing of BC_1-2_ (Fig. 3C, Fig. 4). Comparison of these genotypes showed that, as expected for plants backcrossed to Optic, all chromosomes contained one or two discrete segments of the M1460 genome in an otherwise Optic genome background. However, only two introgressed regions were conserved in all of the transgenic plants. The first region is on chromosome 5H and includes the *LYS*3 locus. Conservation of this region is expected since the shrunken grain phenotype typical of *lys3* mutants was used to select the BC lines. At least some of the selected lines are heterozygous at the *LYS3* locus suggesting that selection for homozygous *lys3* genes based on shrunken-endosperm phenotype was not perfect. That some of the BC_3_ and BC_4_ generations possess the *lys3* gene but are not transformable suggests that transformability is not due to the *lys3* gene (or to any other gene in the 5H-conserved region).

**Figure 4.**
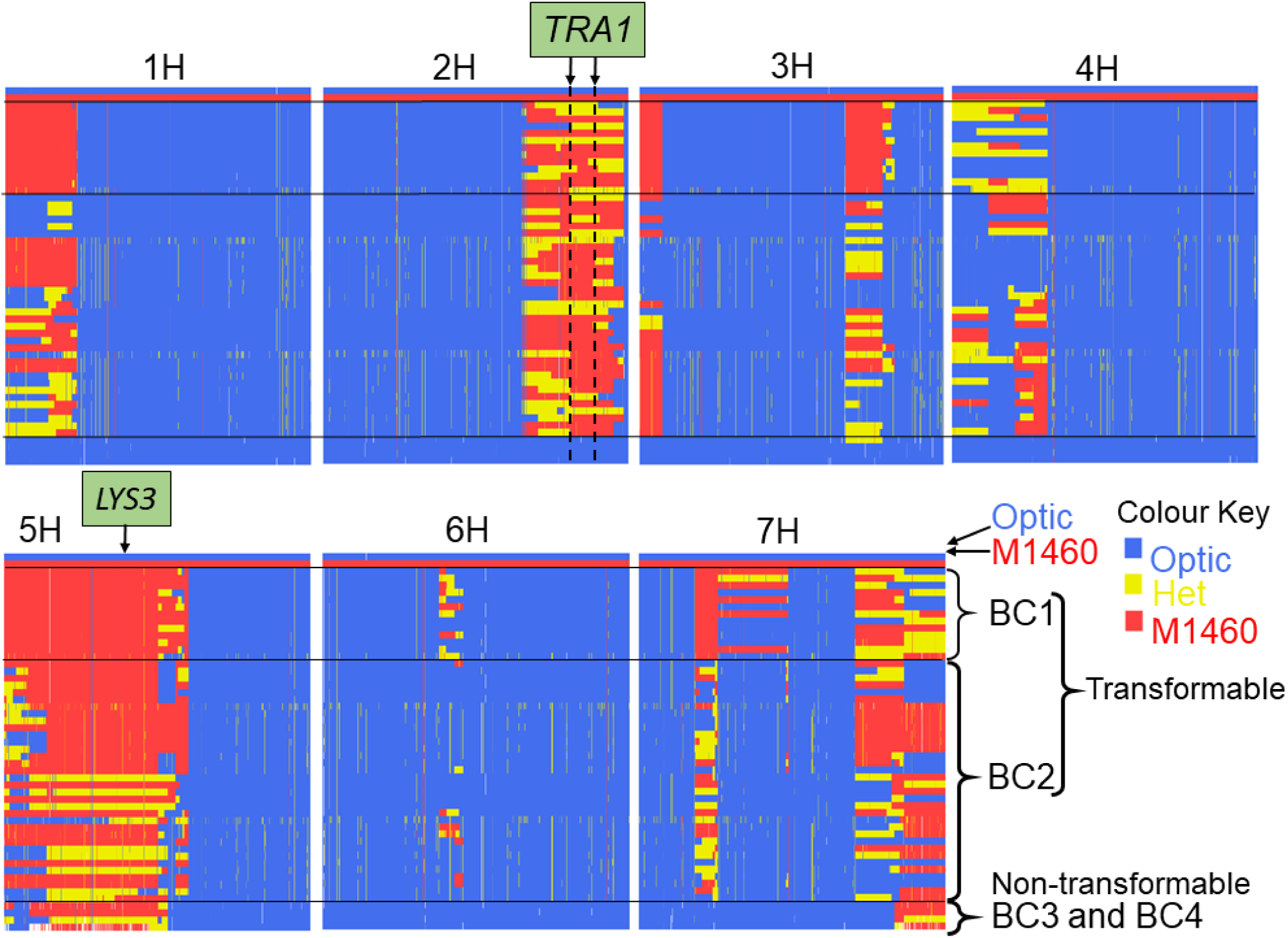
Comparison of the genotypes of transformable and non-transformable plants. The genotypes of 13 BC1 transgenic BC1 plants, 35 BC2 transgenic BC2 plants and four non-transformable embryo-donor BC3 and BC4 plants were compared using the Barley 50k iSelect SNP array. All BC1 and BC2 embryo donor plants were transformable, whereas all of the BC3 and BC4 embryo-donor plants were recalcitrant to transformation. Polymorphic markers are in columns arranged according to their physical position in the 7 chromosomes of barley. Blue = marker with Optic genotype, red = M1460 genotype and yellow = heterozygous genotype. The positions of the *LYS3* gene and the *TRA1* region are indicated. SNP genotype data are provided in Supplementary Table 1.

The second conserved region is on chromosome 2H and it is conserved in the 48 transgenic BC_1_ and BC_2_ plants but not in the four non-transformable BC_3_ and BC_4_ embryo-donor plants (Fig. 4 and Fig 5A). These data suggest that within this region on chromosome 2H, there is a transformability gene (or genes) inherited from M1460. Although the region is conserved, some or all genes in the transformed plants are heterozygous suggesting that the transformability gene may be dominant. We named this locus *TRA1* (*Transformability factor 1*). In the reference cv. Morex, the *TRA1* region is 11.2 Mbp and contains 225 high-confidence genes (RefSeqv2, IBSC) (Fig. 5B).

**Figure 5.**
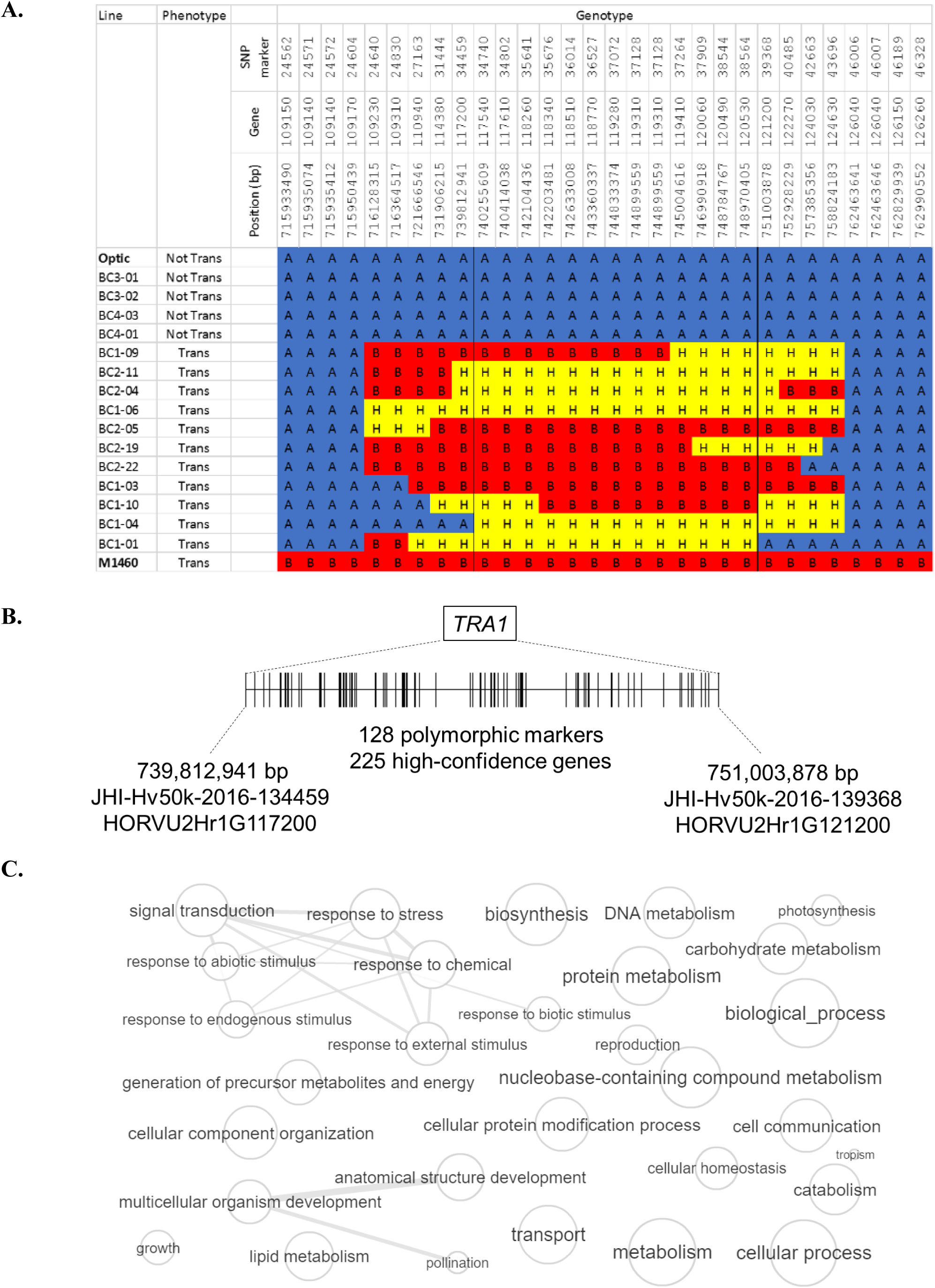
The genomic region containing *TRA1* on chromosome 2H. **A**. Genotype and phenotype data for selected lines from the mapping population (Fig. 4) are shown to illustrate the introgressed region on chromosome 2H that was inherited from M1460. ‘Not Tran’ = not transformable. ‘Trans’ = transformable. Markers with M1460 genotype are shown as ‘B’ and coloured red, Optic are ‘A’ and blue, and heterozygous markers are ‘H’ and yellow. Genotypes of the transformable lines are for plants recovered from transformation assay whilst the genotype of the non-transformable lines are for plants used as embryo donors. All gene names are preceded by: HORVU2Hr1G, whilst the marker names are preceded by JHI-Hv50k-2016-1. The boundaries of the introgressions in plants BC1-04 and BC2-30 define the smallest contribution from M1460 and therefore delimit the *TRA1* region. **B**. A diagram of the *TRA1* region is shown. The two genes indicated correspond to the flanking markers identified by mapping (Fig. 4 and Fig. 5A). Genes (shown as vertical lines) are ordered and positioned according to the barley physical map (Refseqv2, IBSC). **C** The gene categories (GOs) of genes in the *TRA1* region were reduced and visualized using enrichment analysis. Bubble size indicates the frequency of the GO term and highly similar GO terms are linked by edges in the graph (http://revigo.irb.hr/revigo.jsp).

### *TRA1* Candidate Genes

To identify candidate *TRA1* gene(s), first we looked for barley orthologs of genes already known to influence *Agrobacterium*-mediated transformation. The ectopic over-expression of genes involved in cell growth and differentiation has been used to improve transformability in dicotyledonous species (reviewed in Gordon-Kamm *et al*., 2019) and in monocots (Lowe *et al*., 2018). In cereals, the over-expression of either of the genes *WUSCHEL* or *BABY BOOM* stimulates *Agrobacterium*-mediated transformation in maize, rice (*Oryza sativa* L.), sorghum (*Sorghum bicolor* L.) and sugarcane (*Saccharum officinarum* L.) (Lowe *et al*., 2018). However, the barley orthologs of these two genes are not located in the *TRA1* region, or in the QTL regions associated with transformability in Golden Promise (Hisano *et al*., 2017). The *WUSCHEL* ortholog is on chromosome 3H (HORVU3Hr1G085050; chr3H:610834730-610834996) and the *BABY BOOM* orthologue is on chromosome 2H (HORVU2Hr1G087310; chr2H:627137907-627140548).

In *Arabidopsis*, progress has been made in identifying the molecular mechanisms governing *Agrobacterium*-mediated transformation. Although the methods used to transform *Arabidopsis* (floral dipping and root culture) differ from that used to transform barley (embryo culture), we looked to see if any of the *Arabidopsis* transformation genes were *TRA1* candidates. Some of these *Arabidopsis* genes affect somatic embryogenesis but we have not considered these since *TRA1* appears to affect *Agrobacterium* interactions. We therefore considered as *TRA1* candidates 49 *Arabidopsis* genes involved in *Agrobacterium* perception, DNA transfer/integration or tumour development. (Table S1). Mutants affected in these genes, or transgenic plants overexpressing these genes, are resistant or less sensitive, or show increased susceptibility to *Agrobacterium*. However, none of the 49 genes has a predicted orthologue(s) in the *TRA1* region of barley (Table 1).

To identify candidate genes in the *TRA1* region, we sunsequently studied their functional annotations. Analysis of Gene Ontologies (GOs) showed that 141 of the 225 genes have GO annotations and that these are involved in a wide range of biological processes (Fig. 5C). These genes included ones involved in response to stress or signal transduction (GO:0009719, GO:0009628, GO:0009607, GO:0006950, GO:0009605, GO:0007165), cell communication (GO:0007154), growth GO:0048856) or development (GO:0009856, GO:0007275). No transcript data is currently available for *Agrobacterium*-inoculated embryos in barley but an analysis of the response of wheat callus to *Agrobacterium* showed that 24 genes were differentially expressed at the RNA and protein levels (Zhou *et al*., 2013). Of these 24 wheat genes, 17 have 1-to-1 orthologs in barley but none are in *TRA1* region (data not shown).

### The Barley Cultivar Minerva is not the Parent of M1460

As M1460 was reported to have been produced by chemical mutagenesis of the cultivar Minerva (Aastrup, 1983), we compared their genotypes. This unexpectedly revealed that the genotype of M1460 differs significantly (by 14%) from that of Minerva. The level of genotype similarity observed for M1460 and Minerva is within the range typically observed for spring barley cultivars that are not directly related (Malcolm Macauley, personal communication). To investigate this further, ten different accessions of Minerva were ordered from three different germplasm repositories and genotyped using the Barley 50K iSelect SNP array. A phylogenetic analysis of all 11 different Minerva accessions, M1460 and Optic (as a spring barley control) showed that M1460 and Optic are clearly genetically distinct from the ‘Minervas’ (Fig. S4A). Principal Coordinate Analysis (PCA) showed that all 11 Minerva accessions form a cluster, suggesting that they are genuinely closely related, whereas M1460 and Optic occupy two separate positions in PCA space (Fig. S4B). For comparison, we also compared the relatedness of the *lys3* mutants Risø18, Risø19 and Risø1508 and their parent Bomi using the same Principal Coordinate Analysis method (data not shown). This showed the mutants and their parent were closely related. Previously, the use of other genetic markers (RAPDs, AFLPs and SSRs) confirmed the isogenic relationship between Maythorpe and Golden Promise (Forster, 2001). Thus, our data strongly suggest that Minerva is not the parent of M1460.

The origin of M1460 is currently unknown. Comparison of M1460 with 1,000 core barley accessions (Darrier *et al*. 2019) using the Barley 50K iSelect SNP array genotypes (data not shown), failed to identify any barley cultivar as the likely genetic background of M1460. The line most similar to M1460 (92% similarity) was cv. Gerkra (HOR 17444), a Spring 2-row malting barley introduced in The Netherlands in 1969. We also found from this comparison that M1460 was genetically distinct from Golden Promise (20% different).

## DISCUSSION

In this study, we showed that a mutant barley, M1460 has a transformation efficiency comparable with that of the model cultivar for barley transformation, Golden Promise. We identified a locus in M1460, *TRA1*, that is responsible for *Agrobacterium*-mediated genetic transformation. *TRA1* from M1460 was transferred by backcrossing to the cultivar Optic, which is recalcitrant to transformation. Only lines including *TRA1* from M1460 were amenable to transformation. The minimum *TRA1* region identified to date is 11.2 Mbp and contains 225 genes. We are now selecting more homozygous recombinant lines to dissect the *TRA1* region further (Fig. 5A).

Previous studies have suggested a more complicated three-locus source of transformability in Golden Promise (Hisano and Sato, 2016; Hisano *et al*., 2017). Two of the three *TFA* loci previously identified in Golden Promise are located on chromosome 2H and one of these, *TFA3*, could possibly overlap with *TRA1* (Supplementary Fig. S5). However, *TFA1* is thought to be the most important of the three loci in Golden Promise and transformability is thought to be conferred by *TFA3* only in the presence of both *TFA1* and *TFA2* (Hisano *et al*., 2017). The discovery in M1460 of a single locus, capable of conferring transformability increases the potential for the creation of new transformable lines. Further backcrossing of *TRA1* into Optic is underway in our lab to create a readily transformable elite malting barley line suitable for gene functional studies. Since Minerva is not the parent of M1460, we are also attempting to identify the correct pedigree.

When we embarked on this study, our hypothesis was that transformability in M1460 might be due to the *lys3* gene since this mutation is known to affect embryo development (Cook *et al*., 2017, Orman-Ligeza *et al*., 2019). Consequently, we selected the M1460 x Optic backcrossed lines at each generation for the presence of *lys3* (using shrunken endosperm as a phenotypic marker for *lys3*). However, we subsequently discovered that the *lys3* locus was not responsible for transformability. BC_3-4_ plants carrying *lys3* but not *TRA1* were not transformable. With hindsight, selection based on *lys3* was unfortunate. The removal of the *lys3* mutation is desirable since it has a detrimental effect on endosperm size and composition (low starch).

Previous attempts to overcome the cultivar-dependence of transformation efficiency in barley have concentrated on protocol optimization. For example, Murray *et al*. (2004) analysed three different Australian cultivars (Schooner, Chebec and Sloop) and obtained transformation efficiencies of 0.6% compared to 4.4% for Golden Promise by lengthening the culture periods on both regeneration and rooting media compared with the standard protocol for Golden Promise. Hensel *et al*., 2008 combined the anti-oxidative property of L-cysteine (which decreases necrosis) with the use of acetosyringone to improve transformation efficiency with *Agrobacterium*. In these experiments, the efficiency of transformation of Golden Promise was increased to 86.7 % and six of nine barley cultivars previously recalcitrant to transformation were transformable at low frequencies (0.2–7.8%). Further improvement of this protocol has enabled the transformation of more than 20 barley cultivars but, prior to the work reported here, the transformation efficiency of Golden Promise remained higher than that of any other cultivar tested in similar conditions (Marthe *et al*., 2015).

The high transformability of Golden Promise may be due in part to its ability to form callus, shoots and roots in tissue culture. However, other cultivars are equally amenable to tissue culture and this alone is not sufficient to ensure transformability. For example, cv. Morex shows high frequency of plant regeneration from immature embryos but is recalcitrant to transformation (Chang *et al*., *2003;* Hensel *et al*., 2008; our unpublished results). The genotypes used in our experiments were also able to regenerate in tissue culture whether or not they were transformable (cf. Fig. 1C and Fig. 3A). It has been reported that the transformability of Golden Promise may be linked to a higher than normal tolerance to hygromycin selection. Holme *et al*. (2008) found that 23% of the regenerated plants from Golden Promise were escapes (i.e. they grew in the presence of hygromycin but were not transformed with *HPT*). In comparison, all regenerants of the four other cultivars tested by Holme *et al*. (2008) were transgenic and non-chimeric suggesting that they have a lower tolerance to hygromycin than Golden Promise. In contrast, in our transformation system, no escapes were found for M1460 or Golden Promise. We therefore suggest that the efficacy of *TRA1* from M1460 may be due to its ability to interact with *Agrobacterium* and successfully transfer T-DNA. Other barley cultivars that are recalcitrant to transformation may more effectively block *Agrobacterium* infection.

Our work suggests that one (or more) of the 225 genes in the *TRA1* region is responsible for M1460 transformability. Whilst fine mapping is underway to dissect the *TRA1* region further, we attempted to identify candidate genes by studying their annotations and patterns of expression. The positions of orthologs of genes known to affect transformation efficiency in other species was determined but none are located in the *TRA1* region. Examination of the known functions of the genes in the *TRA1* region (analysis of Gene Ontologies) revealed potential candidate genes involved in a range of processes including plant-pathogen response. This is interesting because the process of *Agrobacterium*-mediated transformation essentially mimics the successful infection of a host plant by a pathogen. The mechanisms involved include attachment and recognition between *Agrobacterium* and host plant, production of T-DNA in the bacterium and its transfer to the host cells, integration of the T-DNA into the host genome and finally, its expression. Consistent with this, pathogen-response and/or stress-related genes have been shown in several species to be involved in determining transformability. For example, an *Arabidopsis* mutant with constitutive expression of plant defence-related genes had reduced susceptibility to *Agrobacterium* infection (Veena *et al*., 2003). Also, transfer-competent, but not transfer-incompetent, *Agrobacterium* strains suppress plant defence genes during infection in *Arabidopsis* (Veena *et al*. 2003). In wheat, there have been attempts to identify the genes responsible for transformability by comparing gene expression in embryos cultured in the presence or absence of *Agrobacterium* (Zhou *et al*., 2013). A set of twenty-four genes were found to be differentially expressed at both the RNA and protein levels and most of these genes had roles in stress or immunity-response. Further work is underway to identify the *TRA1* gene in barley. Once identified, its efficacy in a range of barley cultivars (and other species) that are currently recalcitrant to transformation will be tested.

## AUTHOR CONTRIBUTIONS

BOL carried out the bulk of the experimental work. KT and BOL designed the project, with advice from CU and WH. AH performed the tissue culture, plant transformation and husbandry, and WH supervised this work. PH was responsible for the SNP genotyping and together with MM, provided advice to BOL on its analysis. BOL and KT wrote the paper together and all authors proof-read the paper and provided feedback. Authors, apart from the first and last, are listed alphabetically.

## ACKNOWLEDGMENTS

The authors thank the BBSRC for funding this work. Dr Nicole Schatlowski, NIAB (now EMBL-EBI, Wellcome Genome Campus, Hinxton, Cambs, UK) is thanked for her contribution to the Optic-M1460 backcrossing programme. Dr Cong Tan, The James Hutton Institute, Invergowrie, Dundee DD2 5DA, Scotland, UK is thanked for preliminary analysis of the SNP genotyping data.

## SUPPLEMENTARY MATERIAL

The Supplementary Material for this article can be found online at: https://www.frontiersin.org/articles/.

**Supplementary File S1. Polymorphic markers (Optic vs M1460) used in this study**.

**Supplementary File S2. Minerva SNP genotypes**.

## Supplementary material

**Supplementary Figure S1.**
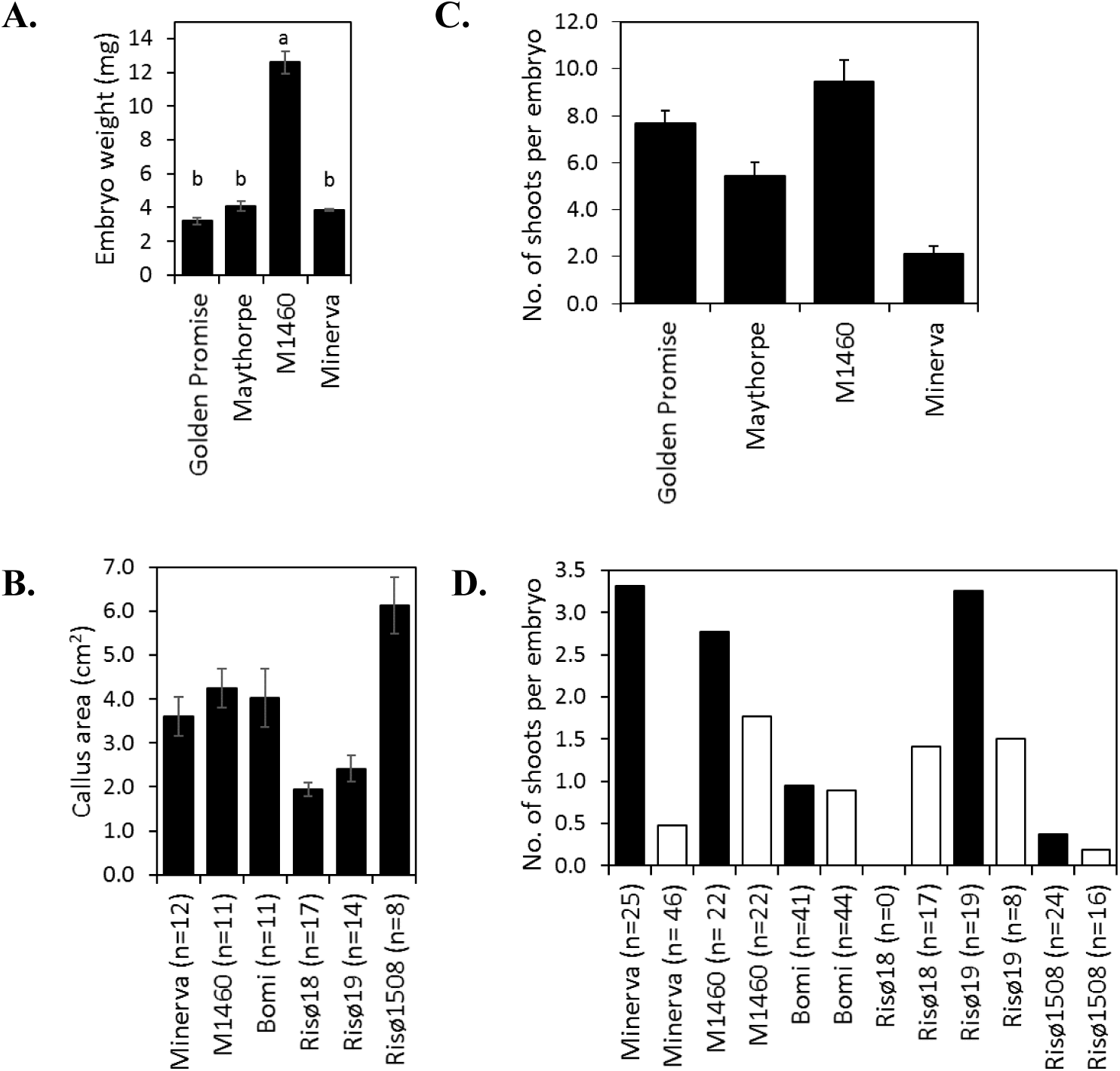
The responses of embryos in tissue culture. A. Embryos were extracted from mature grains and weighed. Maythorpe is the parent of the chemically-induced mutant Golden Promise and Minerva is the parent of the *lys3* mutant M1460. Values are means ± SE for 3 replicate plants (the weight of 10 embryos per plant was determined and the average embryo weight was calculated). Letters indicate statistical significance: data with the same letter are not significantly different (Tukey HSD test, p-value>0.05). B. Embryos (1.5 - 2.0 mm in diameter for Golden Promise and the equivalent developmental stage for the mutant lines) were excised from developing grains, grown on callus induction medium and then on transition medium (both without hygromycin) to induce callus formation. The areas of the resulting calli were measured. Data are means ± SE for calli on two culture plates (n = number of calli). All data are from a single experiment. C. Calli from plates as in (B) were transferred to regeneration medium. The numbers of regenerating shoots (>0.5 cm) per embryo was determined. Values are means ± SE for 35 or 36 embryos from a single experiment. The values for Maythorpe and Minerva were significantly different from Golden Promise (Student’s *t*-test, p-value<0.5). D. Calli from plates as in (B) were transferred to regeneration medium. The numbers of regenerating shoots (>0.5 cm) per embryo was determined. Values for two independent experiments are shown.

**Supplementary Figure S2.**
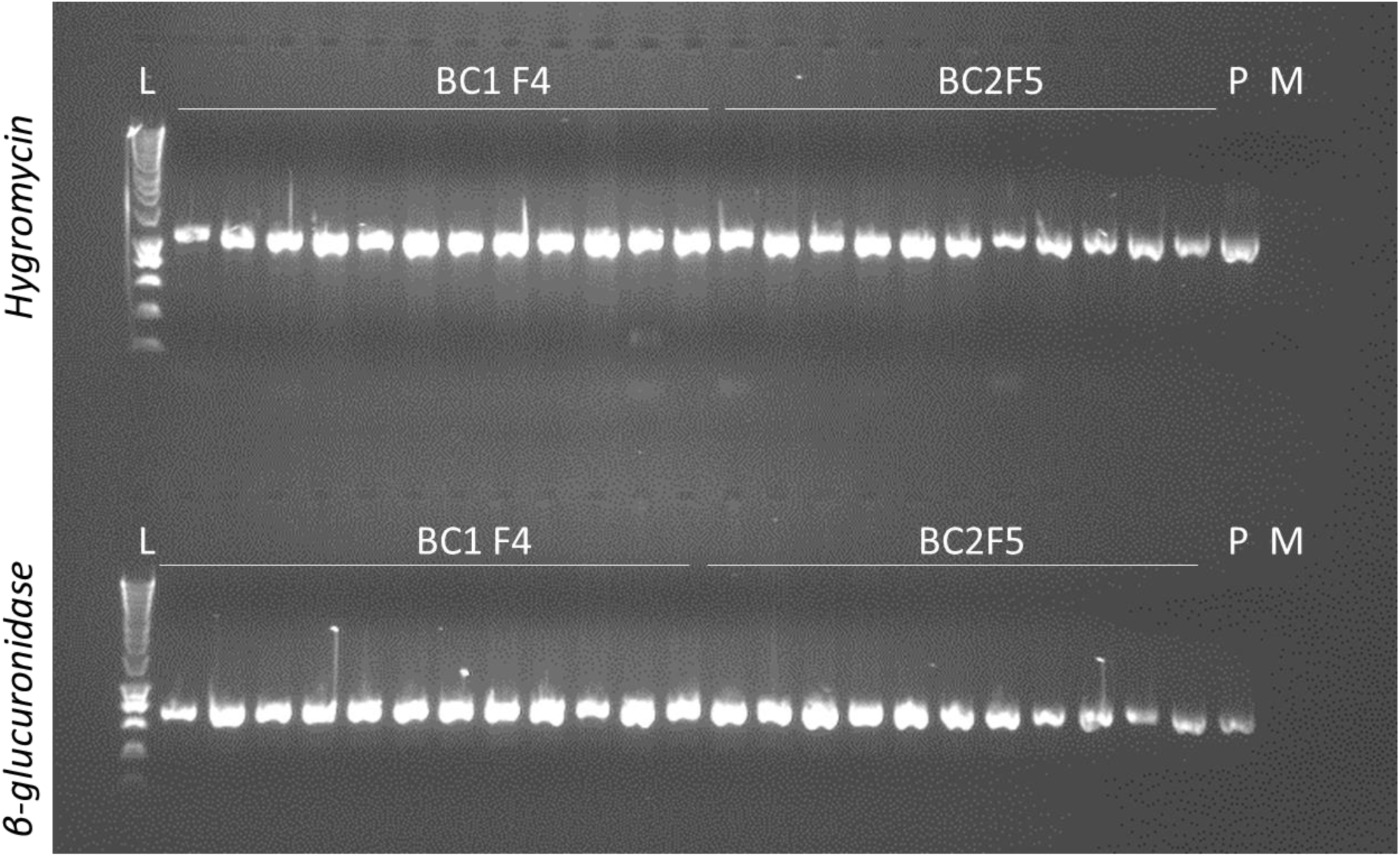
Testing for the presence of transgenes. PCR analysis of transgenic plants. PCR was performed for detecting transgenes using primers specific to hygromycin phosphotransferase (HPT, 1035 bp) and β-glucuronidase (GUS, 710 bp) genes in regenerated plants. L = size standards, P = plasmid positive control where *35S:GUS* in pBract204 was used as a template, M = non-transgenic M1460 plant as negative control.

**Supplementary Figure S3.**
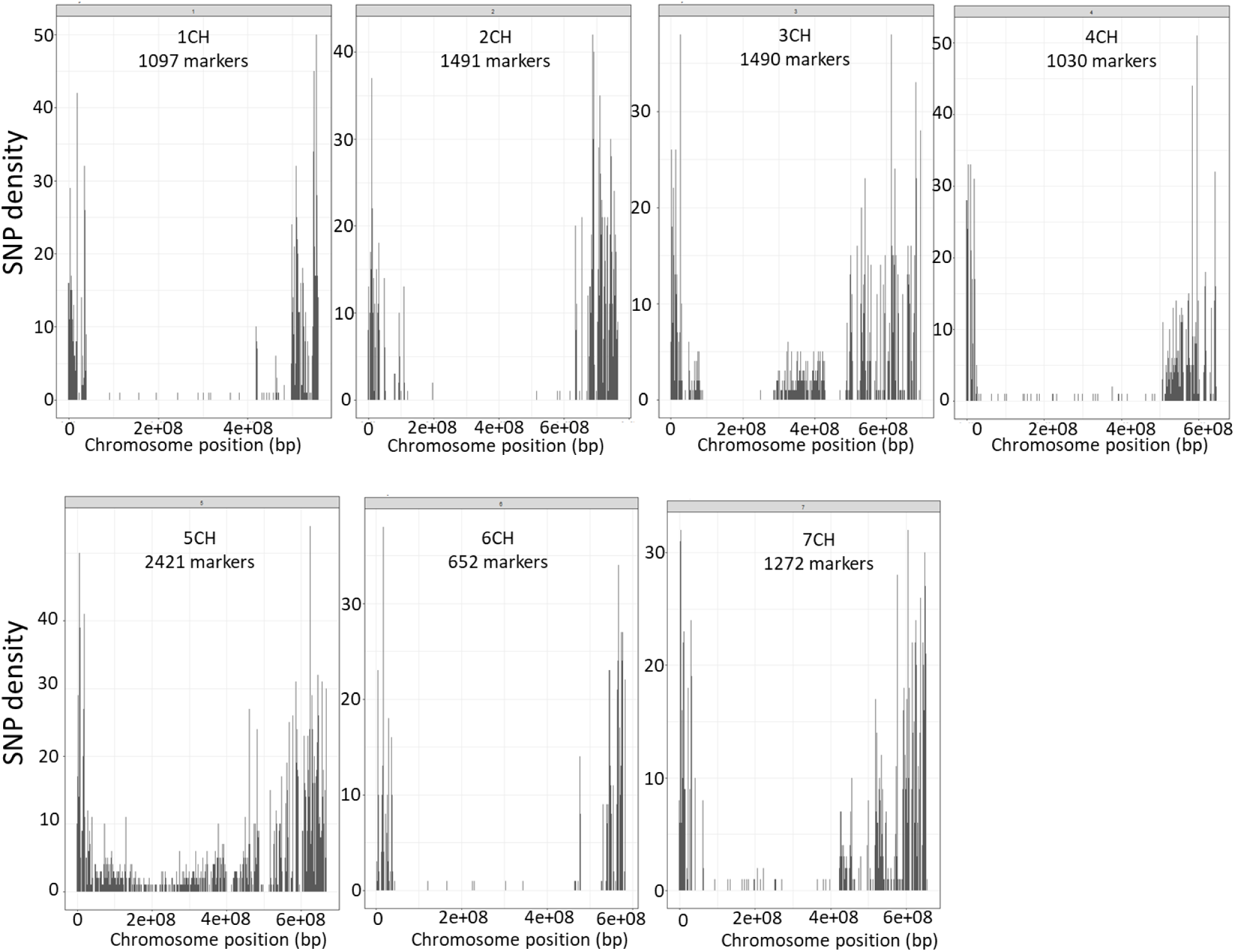
The distribution of polymorphic markers. Genotyping was performed using the Barley 50k iSelect SNP array (Bayer *et al*., 2017). The numbers of polymorphic markers and their distributions across the seven barley chromosomes are shown.

**Supplementary Figure S4.**
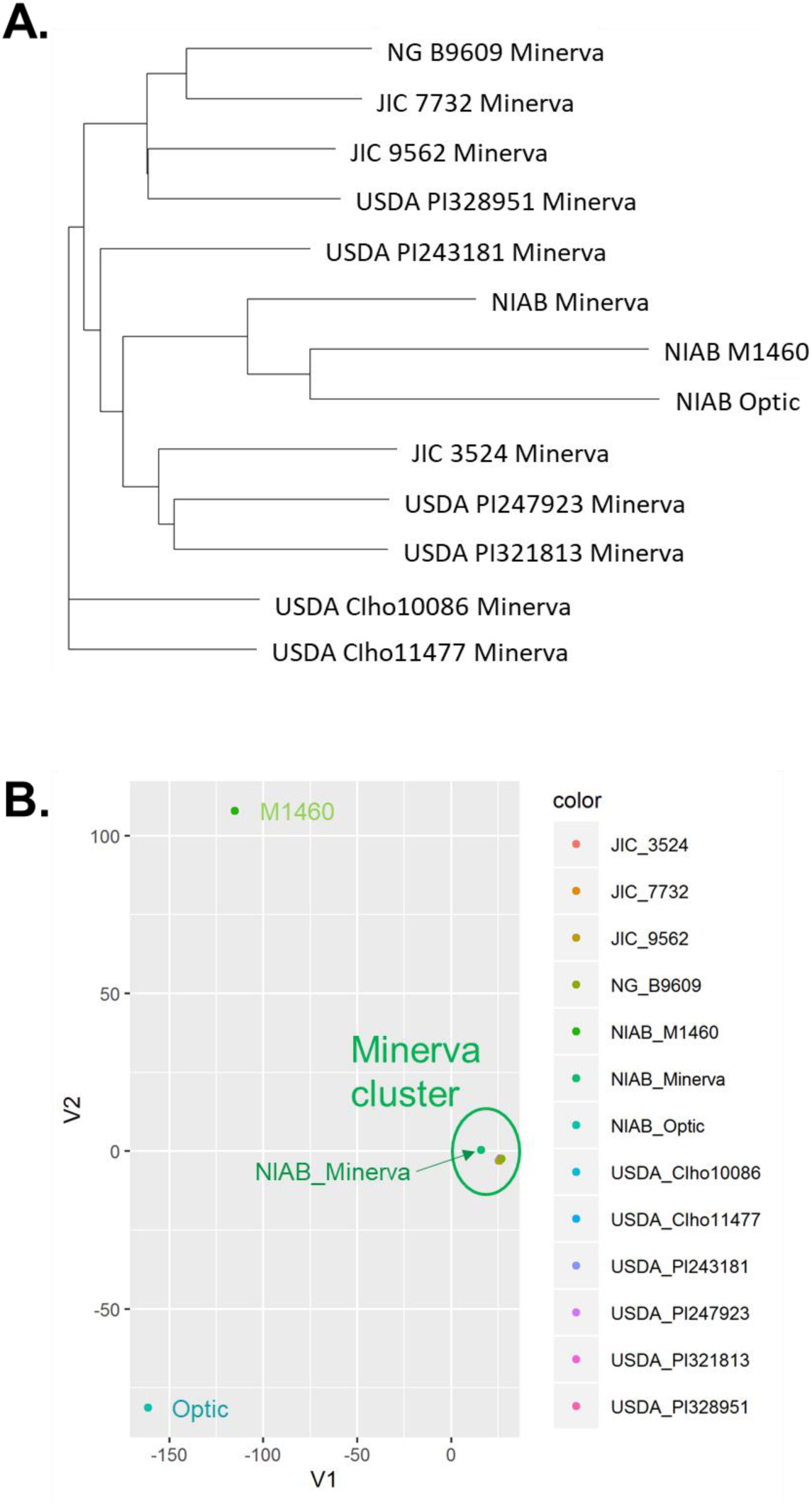
The genotypes of Minerva accessions. To investigate the relationship between M1460 and Minerva, 11 different Minerva accessions, Optic and M1460 were genotyped. SNP genotype data are given in Supplementary File S2. A. An unrooted phylogenetic tree constructed by the neighbour-joining method based on *Euclidean* distance matrix. B. Principal component analysis of genotypes based on *Euclidean* distance matrix.

**Supplementary Figure S5.**
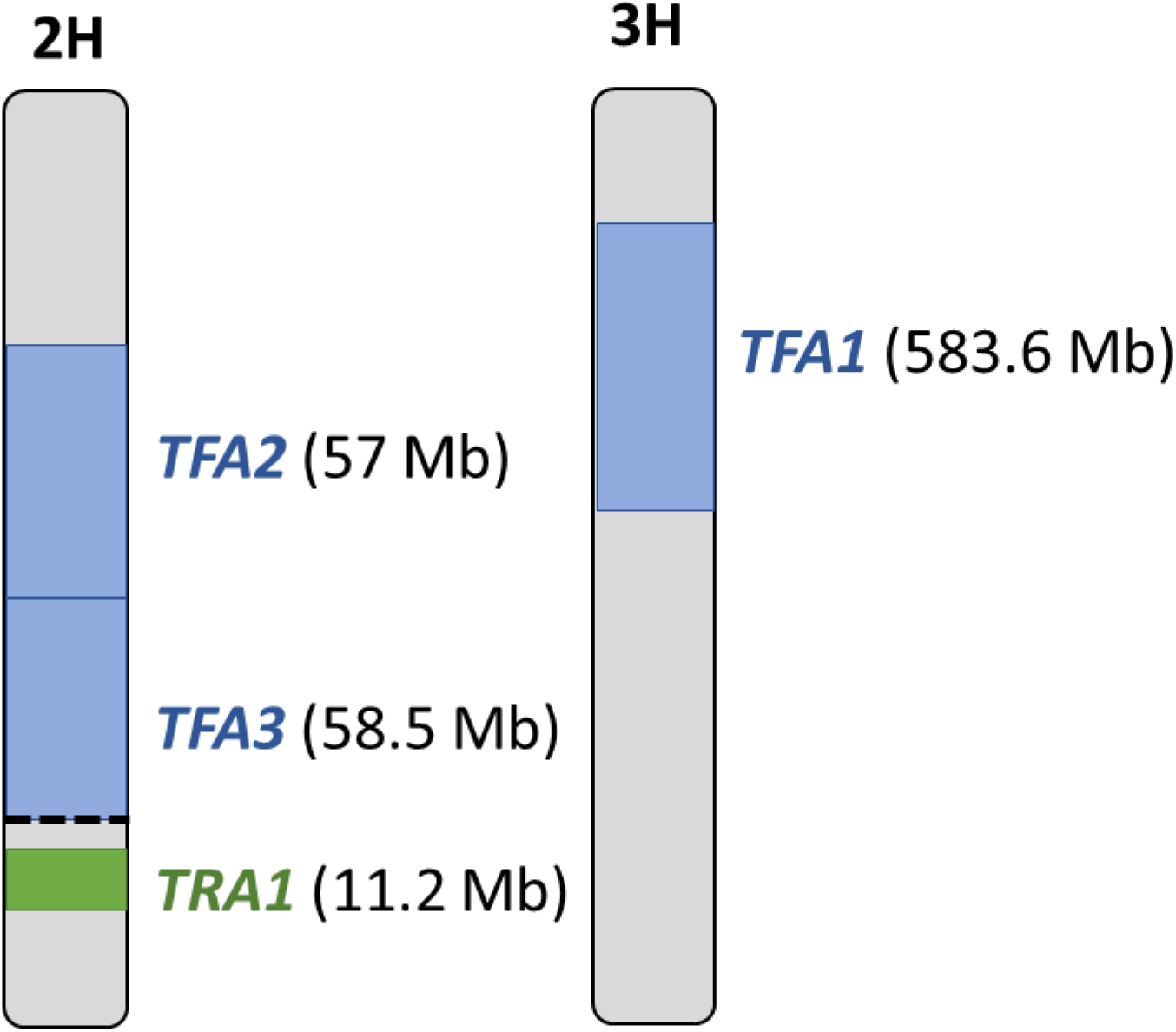
A summary of the transformability regions of barley. Regions of chromosomes 2H and 3H that contain transformability regions are shown and their size is indicated. *TRA1* was identified in M1460 in this study. *TFA1, TFA2* and *TFA3* were identified in Golden Promise by Hisano *et al*. (2017). In M1460, the region that confers transformability lies between markers JHI-Hv50k-2016-134459 (2H: 739812941 bp) and JHI-Hv50k-2016-139368 (2H:751003878 bp). In Golden Promise, *TFA* regions are flanked by the following markers: *TFA1*: NIASHv1109O03_00000798_3H (3H:24,998,400 bp) and BOPA1_8984-579 (3H:608,636,481 pb), *TFA2*: FLOUbaf102l04_00000319_2H (2H:512,036,920 bp) and FLOUbaf138j23_00000441_2H (2H:569,068,240 bp), *TFA3*: FLOUbaf138j23_00000441_2H (2H:569,068,240 bp) and FLOUbaf102a14_00001505_2H (2H:627,567,720 bp, p<0.05). The distal boundary for *TFA3* was not determined with high confidence (see Hisano *et al*., 2017 for details) and is, therefore, shown as a dashed line.

**Supplementary Table S1.** An overview of genes involved in *Agrobacterium*-mediated transformation in *Arabidopsis* and their putative orthologs in barley. This list of *Arabidopsis* genes is adapted from Hwang, Yu and Lai (2018). Putative orthologs in barley were as defined by Ensembl Plants (https://plants.ensembl.org/). The genomic location of the predicted ortholog in barley is shown in brackets for genes on 2H only. References are as given in Hwang, Yu and Lai (2018) with the following additional references: Mysore *et al*., 2000, Zipfel *et al*., 2006, Chinchilla *et al*., 2007, Huang *et al*., 2018.

| *Arabidopsis* gene ID | Function | Putative barley ortholog (Ensembl Plant) | Reference |
| --- | --- | --- | --- |
| Attachment of *Agrobacterium* to plant cells (7 genes) |  |  |  |
| At2g23120 | AGP17 (RAT1, arabinogalactan protein) | Undetermined | Gaspar *et al.*, 2004 |
| At5g03760 | CSLA (RAT4, cellulose synthase-like A9) | HORVU7Hr1G092990<br>HORVU7Hr1G092910 | Zhu *et al.*, 2003 |
| At2g40970 | MTF1 (Myb family transcription factor) | HORVU1Hr1G064950 | Sardesai *et al.*, 2013 |
| At3g28290 | AT14A (similar to integrins) | HORVU2Hr1G006810<br>(chr2H:14,214,133-14,214,948)<br>HORVU4Hr1G070530 | Sardesai *et al.*, 2013 |
| At5g20480 | EFR (EF-Tu receptor that is required for perception of the bacterial PAMP EF-Tu) | Undetermined | Zipfel *et al.*, 2006 |
| AT5G46330 | FLS2 (leucine-rich repeat receptor kinases flagellin-sensitive 2) | HORVU2Hr1G104030<br>(chr2H:701,642,507-701,647,785) | Chinchilla *et al.*, 2007 |
| AT4G33430 | BAK1 (BRI1-associated receptor kinase 1) | Undetermined | Chinchilla *et al.*, 2007 |
| Transport of virulence proteins (6 genes) |  |  |  |
| At4g23630 | RTNLB1 (reticulon-like protein B-1) | Undetermined | Hwang and Gelvin, 2004 |
| At4g11220 | RTNLB2 (reticulon-like protein B-2) | Undetermined | Hwang and Gelvin, 2004 |
| At5g41600 | RTNLB4 (reticulon-like protein B-4) | Undetermined | Hwang and Gelvin, 2004 |
| AT1G64090 | RTNLB3 (reticulon-like protein B-3) | Undetermined | Huang *et al.*, 2018 |
| AT3G10260 | RTNLB8 (reticulon-like protein B-8) | HORVU2Hr1G060770<br>(chr2H:406,063,544-406,071,319)<br>HORVU5Hr1G101680 | Huang *et al.*, 2018 |
| At3g53610 | RAB8B (small GTPase) | HORVU1Hr1G072070<br>HORVU3Hr1G086830<br>HORVU5Hr1G112790 | Hwang and Gelvin, 2004 |
| Import of T-DNA and effector proteins (11 genes) |  |  |  |
| At4g38740 | ROC1 (cyclophilin proteins, peptidyl-prolyl cis-trans isomerase CYP1) | HORVU6Hr1G012570<br>HORVU6Hr1G092950 | Deng *et al.*, 1998 |
| At4g26080 | PP2C (type 2C protein phosphatase, ABI1) | HORVU1Hr1G080290<br>HORVU1Hr1G094840<br>HORVU3Hr1G050340<br>HORVU3Hr1G067380<br>HORVU7Hr1G029040 | Tao *et al.*, 2004 |
| At3g06720 | KAPα/IMPA-1 (Karyopherin protein, importin alpha isoform 1 involved in nuclear import) | Undetermined | Bhattacharjee *et al.*, 2008 |
| At4g16143 | IMPA-2 (importin alpha isoform 2) | Undetermined | Bhattacharjee *et al.*, 2008 |
| At4g02150 | IMPA-3 (importin alpha isoform 3) | HORVU3Hr1G034750 | Bhattacharjee *et al.*, 2008 |
| At1g09270 | IMPA-4 (importin alpha isoform 4) | HORVU2Hr1G016300 | Bhattacharjee *et al.*, 2008 |
| At1g43700 | VIP1 (VirE2-interacting plant protein 1) | HORVU4Hr1G020540<br>HORVU5Hr1G039870 | Tzfira *et al.*, 2002 |
| At5g59710 | VIP2 (VirE2-interacting plant protein 2) | HORVU1Hr1G069060<br>HORVU1Hr1G085740<br>HORVU5Hr1G077990<br>HORVU5Hr1G078050<br>HORVU6Hr1G095320 | Tzfira *et al.*, 2002 |
| At3g45640 | MPK3 (mitogen-activated protein kinase 3) | HORVU4Hr1G057200 | Djamei *et al.*, 2007 |
| At3g18780 | Microfilament and ACT2 (actin gene) | Undetermined | Zhu *et al.*, 2003<br>Yang *et al.*, 2017 |
| At5g09810 | ACT7 (actin gene) | HORVU1Hr1G002840<br>HORVU1Hr1G074350<br>HORVU5Hr1G039850<br>HORVU5Hr1G117900 | Zhu *et al.*, 2003 |
| Integration and/or expression of T-DNA (25 genes) |  |  |  |
| At5g57160 | LIG4 (DNA ligase IV) | HORVU2Hr1G100690 (chr2H:691,977,835-691,978,386)<br>HORVU2Hr1G100740 (chr2H:692,063,686-692,072,163) | Friesner and Britt, 2003<br>van Attikum *et al.*, 2003<br>Mestiri *et al.*, 2014<br>Park *et al.*, 2015 |
| At1g48050 | KU80 | HORVU0Hr1G038620 | Friesner and Britt, 2003<br>van Attikum *et al.*, 2003<br>Mestiri *et al.*, 2014<br>Park *et al.*, 2015 |
| At1g16970 | KU70 | HORVU5Hr1G012090 | Friesner and Britt, 2003<br>van Attikum *et al.*, 2003<br>Mestiri *et al.*, 2014<br>Park *et al.*, 2015 |
| At5g54260 | MRE11 (meiotic recombination 11) | HORVU2Hr1G116540 (chr2H:738,323,340-738,341,229)<br>HORVU7Hr1G085450 | van Attikum *et al.*, 2003 |
| At1g80420 | XRCC1 (homolog of X-ray repair cross complementing 1) | HORVU7Hr1G019990 | Mestiri *et al.*, 2014<br>Park *et al.*, 2015 |
| At5g64520 | XRCC2 (homolog of X-ray repair cross complementing 2) | HORVU3Hr1G082620 | Mestiri *et al.*, 2014<br>Park *et al.*, 2015 |
| At3g23100 | XRCC4 (homolog of X-ray repair cross complementing 4) | Undetermined | Vaghchhipawala *et al.*, 2012<br>Park *et al.*, 2015 |
| At5g41150 | XPF/RAD1/UVH1 (ultraviolet hypersensitive 1) | HORVU4Hr1G090160<br>HORVU4Hr1G090210 | Nam *et al.*, 1998<br>Mestiri *et al.*, 2014 |
| At2g31320 | PARP1 (poly(ADP-ribose) polymerases 1) | HORVU1Hr1G005980 | Jia *et al.*, 2012<br>Park *et al.*, 2015 |
| At5g54640 | HTA1 (histone H2A) | HORVU6Hr1G092280<br>HORVU6Hr1G092390<br>HORVU7Hr1G112470 | Tenea *et al.*, 2009<br>Mysore *et al.*, 2000 |
| At1g43700 | VIP1 (VirE2-interacting plant protein 1) | HORVU4Hr1G020540<br>HORVU5Hr1G039870 | Tzfira *et al.*, 2001<br>Tzfira *et al.*, 2002 |
| At5g59710 | VIP2 (VirE2-interacting plant protein 2) | HORVU1Hr1G069060<br>HORVU1Hr1G085740<br>HORVU5Hr1G077990<br>HORVU5Hr1G078050<br>HORVU6Hr1G095320 | Tzfira *et al.*, 2002<br>Anand *et al.*, 2007 |
| At4g38130 | HDA1/HDA19 (histone deacetylase 1/ histone deacetylase 19) | HORVU6Hr1G038710<br>HORVU7Hr1G085870 | Zhu *et al.*, 2003<br>Crane and Gelvin, 2007<br>Gelvin and Kim, 2007 |
| At3g44750 | HDA2/HDT1 (histone deacetylase 2A) | HORVU1Hr1G095140 | Zhu *et al.*, 2003<br>Crane and Gelvin, 2007<br>Gelvin and Kim, 2007 |
| At5g22650 | HD2B/HDT2 (histone deacetylase 2B) | Undetermined | Zhu *et al.*, 2003<br>Crane and Gelvin, 2007<br>Gelvin and Kim, 2007 |
| At5g38110 | SGA1/ASF1B (anti-silencing function 1B) | HORVU1Hr1G084120<br>HORVU3Hr1G063450 | Gelvin and Kim, 2007 |
| At1g65470 | FAS1 (fasciata 1/nucleosome/ chromatin assembly factor group B/ chromatin assembly factor-1 (CAF-1) subunit) | HORVU5Hr1G021430<br>HORVU5Hr1G084610 | Endo *et al.*, 2006 |
| At5g64630 | FAS2 (fasciata 2/nucleosome/chromatin assembly factor group B/chromatin assembly factor-1 (CAF-1) subunit) | HORVU7Hr1G077320 | Endo *et al.*, 2006 |
| At1g75950 | ASK1/SKP1 (SKP1 homologue 1) | Undetermined | Schrammeijer *et al.*, 2001<br>Tzfira *et al.*, 2002<br>Zaltsman *et al.*, 2010<br>Anad *et al.*, 2012 |
| At5g42190 | ASK2 (SKP-like 2) | HORVU5Hr1G092210 | Schrammeijer *et al.*, 2001<br>Tzfira *et al.*, 2002, 2004<br>Zaltsman *et al.*, 2010<br>Anad *et al.*, 2012 |
| At3g21860 | ASK10 (SKP-like 10) | Undetermined | Schrammeijer *et al.*, 2001<br>Tzfira *et al.*, 2002, 2004<br>Zaltsman *et al.*, 2010<br>Anad *et al.*, 2012 |
| At4g23570 | SGT1a (suppressor of the G2 allele of SKP1) | HORVU3Hr1G055920 | Anand *et al.*, 2012 |
| At4g11260 | SGT1b/EDM1 (Enhanced downy mildew 1) | HORVU3Hr1G055920 | Anand *et al.*, 2012 |
| At1g56250 | VBF (VIP1-binding F-box protein) | HORVU0Hr1G021610<br>HORVU2Hr1G104500 (chr2H:703,924,767-703,927,102)<br>HORVU3Hr1G014270<br>HORVU5Hr1G015260<br>HORVU5Hr1G124090<br>HORVU6Hr1G089430<br>HORVU6Hr1G089480<br>HORVU6Hr1G090000<br>HORVU6Hr1G090010<br>HORVU7Hr1G012610<br>HORVU7Hr1G072570 | Niu *et al.*, 2015 |
| At4g32700 | TEBICHI/POLQ (DNA polymerase theta, Pol $\theta$) | HORVU0Hr1G026710<br>HORVU1Hr1G065680 | van Kregten *et al.*, 2016 |

